# Signaling by a tyrosine-sulfated peptide balances growth and osmotic stress response in rice

**DOI:** 10.64898/2026.08.04.742889

**Authors:** Yejin Shim, Ellen Y. Rim, Jin C.-Y. Liao, Myeong-Je Cho, George Austin, Patrick W. Carlos, Sameer S. Kulkarni, Richard J. Payne, Maria Florencia Ercoli, Pamela C. Ronald

## Abstract

Peptide hormone signaling coordinates plant growth and osmotic stress responses, yet how the transition between these responses is regulated remains poorly understood. Here, we investigated the function of the rice PLANT PEPTIDES CONTAINING SULFATED TYROSINE 8 (OsPSY8) peptide in osmotic stress responses. *OsPSY8* was predominantly expressed in root tissues under non-stress conditions, with preferential expression in lateral roots where it promoted root growth. Osmotic stress rapidly reduced *OsPSY8* expression in roots through the OsWRKY24 transcription factor. Loss-of-function *ospsy8* mutants exhibited enhanced osmotic stress tolerance, whereas *OsPSY8* overexpression increased osmotic stress susceptibility. Transcriptomic analyses revealed that disruption of *OsPSY8* activated stress-responsive pathways, including those associated with lignin biosynthesis, compatible solute production, cell wall remodeling, and reactive oxygen species (ROS) scavenging, and was accompanied by increased lignin accumulation in roots. In contrast, overexpression of *OsPSY8* resulted in maintenance of growth-associated transcriptional programs while suppressing stress-responsive pathways under osmotic stress. Together, these findings identify OsPSY8 as an important regulator of the transition from growth to stress adaptation in rice and suggest that stress-induced repression of PSY signaling is required to disengage growth programs and activate adaptive responses during osmotic stress.

**Significance Statement:** Crop survival during drought depends on the ability to transition from growth to stress adaptation. Plant peptide hormones have emerged as important regulators of this critical transition, highlighting the importance of investigating their roles and potential for improving crop resilience. We show that a rice peptide hormone regulates this transition. Under non-stress conditions, this peptide hormone, predominantly expressed in rice roots, promotes root growth while suppressing stress responses. During osmotic stress, expression of the peptide hormone decreases, resulting in activation of stress-responsive pathways, such as lignin biosynthesis and reactive oxygen species scavenging. These findings demonstrate that a peptide hormone coordinates the balance between growth and stress adaptation in rice, with broader implications for understanding and improving crop resilience.

## Introduction

Plants, as sessile organisms, are continuously exposed to adverse environmental conditions, including extreme temperatures and limited water availability. Among abiotic stressors, drought poses a major threat to crop productivity by compromising photosynthesis, nutrient transport, and grain filling (1). Understanding how plants perceive and respond to changes in water availability is therefore essential for mitigating drought-induced yield losses. Upon drought perception, plants rapidly prioritize survival over growth through the activation of stress-responsive signaling networks and adaptive physiological programs that facilitate acclimation to water-limiting conditions (2).

Recent studies have highlighted small, secreted peptide hormones as important regulators of plant responses to drought stress, mediating the perception of water-deficit signals and the activation of downstream physiological and developmental adaptations (3). These peptide hormones are encoded as precursor proteins, typically composed of fewer than 150 amino acids, that undergo proteolytic processing and, in many cases, post-translational modifications to generate shorter bioactive peptides, often approximately 20 amino acids long (4, 5).hese mature peptides are secreted into the extracellular space and perceived by plasma membrane-localized receptor-like kinases (RLKs), thereby initiating signaling pathways that regulate plant growth and stress responses (4, 5). For example, the expression of the peptide hormone *CLAVATA3/EMBRYO SURROUNDING REGION 25* (*CLE25*) is induced in Arabidopsis root tissues within a few hours in response to dehydration stress (6). The root-derived CLE25 peptide is subsequently transported to leaves through vascular tissues, where it promotes stomatal closure by increasing abscisic acid (ABA) accumulation (6). The leaf-expressed CLE5 peptide acts as a local signal that promotes even faster stomatal closure and drought tolerance independently of ABA accumulation (7). In wheat, the root-derived CLE peptide TaCLE24b promotes lateral root development and enhances drought tolerance (8). Peptide hormones have also been implicated in developmental adaptations to drought stress. For example, *phytosulfokine* (*PSK*) is expressed in tomato abscission zones and promotes drought-induced flower abscission through induction of cell wall hydrolases (9). PSY peptides belong to a class of post-translationally modified peptides whose precursors undergo tyrosine sulfation, a modification required for full biological activity (4). The first PSY peptide was isolated from *Arabidopsis* cell suspension cultures and shown to promote cellular proliferation and expansion (10). In rice, eight *OsPSY* genes have been identified, and OsPSY1, OsPSY2, and OsPSY5 have been reported to promote primary root growth. (11, 12, 13). Beyond growth promotion, PSY signaling has been implicated in coordinating the trade-off between growth and stress responses in plants. In Arabidopsis, exogenous application of the PSY5 peptide promotes root elongation, and loss of its cognate receptors*, PSYR1, PSYR2, and PSYR3* (the *psyr1,2,3* triple mutant), phenocopies this response, demonstrating that the PSY receptors (PSYR) act as negative regulators of growth and that PSY peptides promote growth by repressing PSYR signaling (14). The transcriptome of PSY-treated plants substantially overlaps with that of the *psyr1,2,3* mutant, both characterized predominantly by downregulation of stress-responsive transcription factors, suggesting that a constitutive PSY response represses PSYR-induced stress responses (14). Consistent with this molecular observation, either loss of PSYR or exogenous treatment with the PSY5 peptide compromises plant tolerance to both biotic and abiotic stresses (14). Together, these findings establish that PSY peptides play a central role in the growth-stress response trade-off, whereby PSY peptides repress PSYR-mediated stress responses, whereas the absence of PSY relieves this repression, activating both PSYR-mediated growth restriction and stress responses. In rice, several *OsPSY* genes exhibit altered expression in response to various abiotic stresses, including drought (15), suggesting their potential involvement in stress adaptation.

Despite these findings, the functions of OsPSY peptides in stress responses in rice remain poorly understood. In this study, we characterized the role of OsPSY8 in osmotic stress responses. *OsPSY8* is predominantly expressed in root tissues under non-stress conditions, whereas osmotic stress represses its expression. Overexpression of *OsPSY8* promoted root elongation but increased susceptibility to osmotic stress, whereas loss of *OsPSY8* function enhanced osmotic stress tolerance. Transcriptomic analyses showed that constitutive *OsPSY8* expression promotes growth-associated transcriptional programs and suppresses stress responses, whereas disruption of *OsPSY8* activates stress-responsive pathways, such as lignin biosynthesis. Together, these results suggest that OsPSY8 regulates the transition from growth to stress adaptation and provide insight into peptide-mediated regulation of osmotic stress responses in rice.

## Results

### *OsPSY5* and *OsPSY8* are predominantly expressed in the root and downregulated by osmotic stress

To identify candidate *OsPSY* genes involved in root responses to water availability, we analyzed the expression of all eight *OsPSY* family members in leaves and roots of 9-day-old hydroponically grown rice seedlings by quantitative reverse transcription PCR (qRT-PCR). Among the eight *OsPSY* genes, *OsPSY5* showed the highest expression in roots, followed by *OsPSY4*, *OsPSY8*, and *OsPSY6* (Fig. 1A). Notably, *OsPSY5* and *OsPSY8* transcripts were highly enriched in roots and were barely detectable in leaves, in contrast to *OsPSY4* and *OsPSY6*, which were expressed in both tissues. Histochemical β-glucuronidase (GUS) staining further revealed distinct spatial expression patterns, with *OsPSY5* expressed predominantly in the root stele (Fig. S1A) and *OsPSY8* expressed preferentially in lateral roots (Fig. 1B).

**Figure 1.**
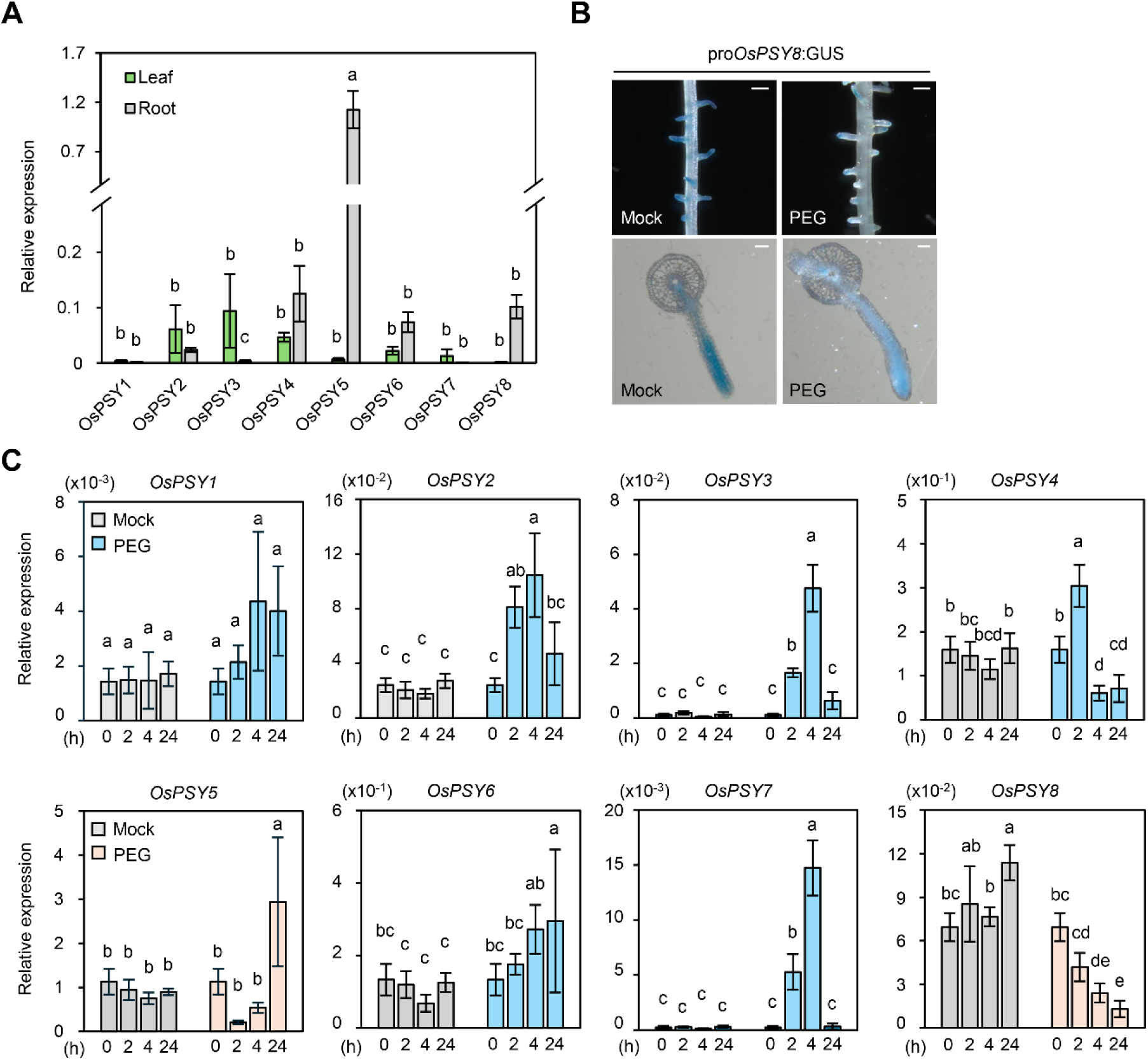
*OsPSY8* is preferentially expressed in roots and repressed by osmotic stress. (A) Spatial expression of eight *OsPSY* genes in leaves and roots of 9-d-old hydroponically grown rice seedlings. Data represent mean ± SD (n = 3 biological replicates), each replicate consisting of three plants. (B) Histochemical GUS staining of *pOsPSY8*::GUS transgenic rice seedlings. Mock, nontreated control; PEG, seedlings treated with 25 % PEG6000 for 4 h. Upper panels, root differentiation zone images. Scale bars = 500 µm; lower panels, 90-µm transverse sections prepared by vibratome sectioning. Scale bars = 100 µm. (C) Time-course expression of *OsPSY* genes under osmotic stress. Roots were harvested at 0, 2, 4, and 24 h after treatment with 25 % PEG6000 or mock (nontreated) control. Relative expression was quantified by RT-qPCR as the transcript level of each target gene, normalized to rice *UBIQUITIN5* (*OsUBQ5*) using the 2^−ΔΔCT^ method. Data represent mean ± SD (n = 3 biological replicates), each replicate consisting of three plants. (A,C) Statistical analyses were conducted using one-way ANOVA followed by Tukey’s honestly significant difference (HSD) post hoc test for multiple comparisons, with different letters indicating significant differences among groups (*P* < 0.05).

We next examined the responses of all eight *OsPSY* genes to osmotic stress using polyethylene glycol (PEG), a well-established, high-molecular-weight, non-penetrating osmotic agent widely used to mimic water-deficit stress in hydroponic systems (16). *OsPSY5* and *OsPSY8* expression decreased as early as 2 h after PEG treatment. By contrast, most other *OsPSY* genes were induced, whereas *OsPSY4* expression increased at 2 h but decreased by 4 h. (Fig. 1C). These trends are consistent with Kesawat et al. (15), where *OsPSY8* (annotated there as *OsPSY4*) was also repressed by osmotic stress in roots, whereas *OsPSY3*, *OsPSY4*, and *OsPSY7*, (annotated there as *OsPSY2*, *OsPSY4*, and *OsPSY7*) were induced. Consistent with the transcript analysis, PEG treatment reduced GUS staining intensity in *pOsPSY8*::GUS transgenic plants compared to untreated controls (Fig. 1B). Together, these results identify *OsPSY5* and *OsPSY8* as the predominant root-expressed members of the *OsPSY* family and show that their expression is repressed in response to osmotic stress.

### Modulation of *OsPSY8* expression altered osmotic stress tolerance in rice

In Arabidopsis, in the presence of PSY peptides, PSYR-mediated stress signaling is suppressed, whereas the absence of PSY peptides activates stress-responsive pathways (14). Given this, we hypothesized that *OsPSY5* and *OsPSY8*, which were repressed under osmotic stress, might function as stress response repressors and selected them for further functional characterization.

To determine whether either peptide regulates osmotic stress responses, we generated *OsPSY5* and *OsPSY8* overexpression lines (*OsPSY5*-OX and *OsPSY8*-OX) under the control of the constitutive *UBIQUITIN* (*Ubi*) promoter and evaluated their performance under PEG-induced osmotic stress. Two independent *OsPSY8*-OX lines (OX1 and OX2) and three independent *OsPSY5*-OX lines (5-OX1, 5-OX2, and 5-OX3) exhibiting high transgene expressions were selected for further analysis (Fig. 2A and S2A). Nine-day-old hydroponic seedlings were exposed to PEG for 4 d and then transferred back to the Hoagland solution for recovery. The survival rate of each genotype was assessed after 7 d of recovery, with plants that remained yellow and desiccated scored as dead and those that produced new leaves scored as surviving. Under these conditions, approximately 40 % of wild-type (WT) plants survived, whereas fewer than 10 % of *OsPSY8*-OX plants recovered (Fig. 2B and 2C). Consistent with this enhanced susceptibility, *OsPSY8*-OX plants exhibited a modest increase in electrolyte leakage during PEG treatment, suggesting greater cellular damage under osmotic stress, although this difference did not reach statistical significance (Fig. 2D). In contrast, *OsPSY5*-OX plants did not show significant differences in survival compared with WT plants under osmotic stress (Fig. S2B and S2C).

**Figure 2.**
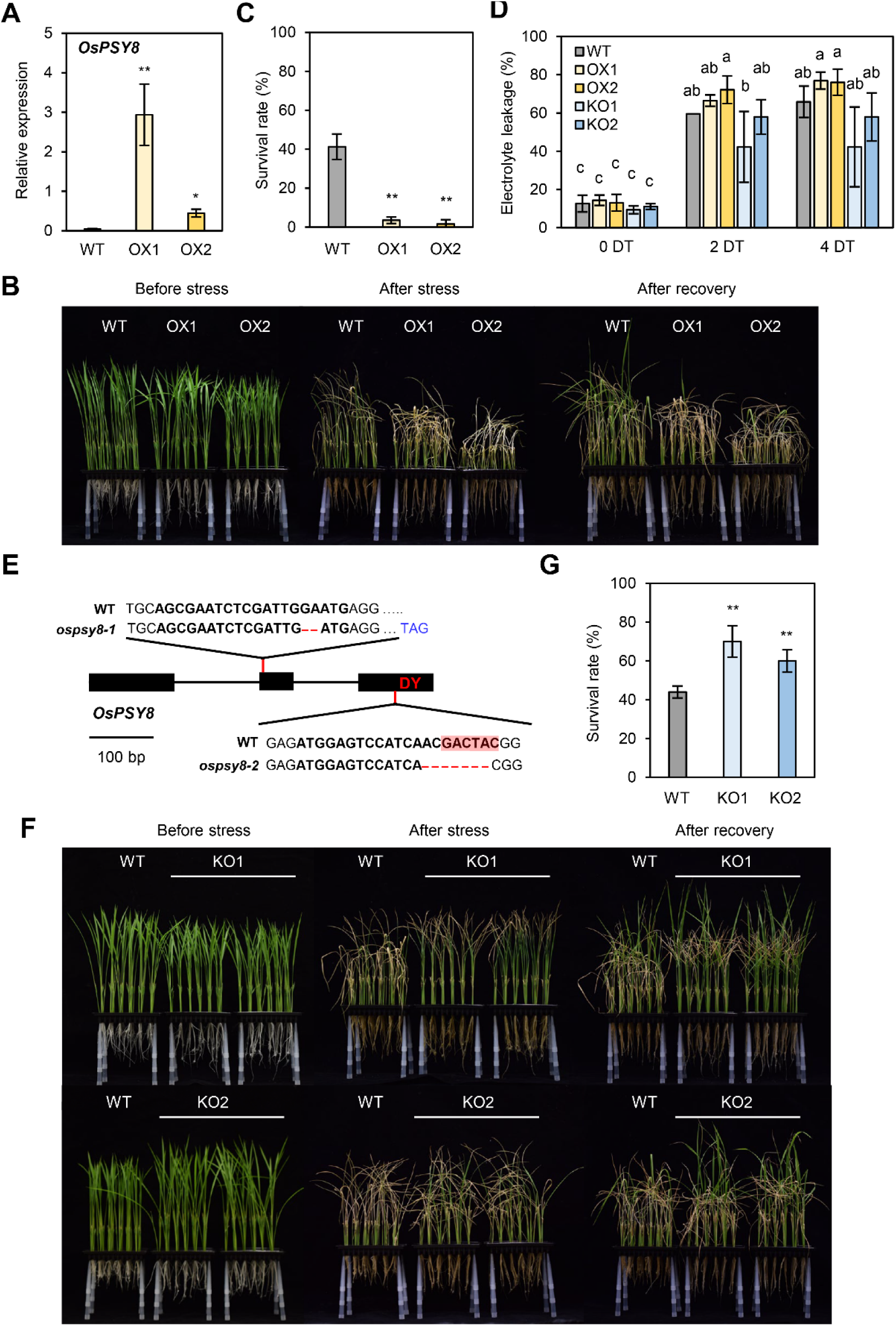
OsPSY8 negatively regulates osmotic stress tolerance in rice. (A) Relative expression levels of *OsPSY8* in WT and two independent *OsPSY8*-OX lines (OX1 and OX2). Relative expression was quantified by RT-qPCR as the transcript level of each target gene, normalized to rice *OsUBQ5* using the 2^−ΔΔCT^ method. (B) Representative photographs of WT, OX1, and OX2 plants before stress, 4 days after PEG-induced osmotic stress, and 7 days after recovery. (C) Survival rates (%) of WT, OX1, and OX2 plants after recovery. (D) Electrolyte leakage (%) measured in WT, OX1, OX2, KO1, and KO2 plants at 0, 2, and 4 days after PEG treatment (DT). Statistical analyses were conducted using one-way ANOVA followed by Tukey’s honestly significant difference (HSD) post hoc test for multiple comparisons, with different letters indicating significant differences among groups (*P* < 0.05). (E) Schematic diagram of the *OsPSY8* gene structure and CRISPR/Cas9-induced mutations. The black boxes represent exons, and the conserved DY motif in the third exon is indicated in red. Sequence alignments show the 2-bp deletion in *ospsy8-1*, which introduces a premature stop codon (TAG, blue), and the 7-bp deletion in *ospsy8-2*, which disrupts the DY motif (red). Dashes indicate deleted nucleotides. (F) Representative photographs of WT, KO1 (*ospsy8-1*), and KO2 (*ospsy8-2*) plants before stress, 4 days after PEG-induced osmotic stress, and 7 days after recovery. (G) Survival rates (%) of WT, KO1, and KO2 plants after recovery. (A, C, G) Data represent mean ± SD (n = 4 biological replicates). Asterisks indicate statistically significant differences from WT (\**P* < 0.05; \*\**P* < 0.01; two-tailed Student’s *t*-test).

We next tested whether loss of *OsPSY8* function would confer increased tolerance to osmotic stress. Two gRNAs targeting the *OsPSY8* coding sequence were used to generate clustered regularly interspaced short palindromic repeats (CRISPR)/CRISPR-associated protein 9 (Cas9) knockout lines. gRNA #1 targeted the second exon, and gRNA #2 targeted the third exon containing the conserved DY motif required for tyrosine sulfation and PSY activity. Target sequences were selected using CRISPRdirect (https://crispr.dbcls.jp/) to minimize potential off-target effects, including targeting of the conserved DY motif in other *OsPSY* genes. Two homozygous mutant alleles were selected for further analysis: *ospsy8-1* (KO1), carrying a 2-bp deletion that introduced a premature stop codon, and *ospsy8-2* (KO2), carrying a 7-bp deletion that disrupted the DY motif (Fig. 2E). Both *ospsy8* mutants displayed enhanced tolerance to osmotic stress. Survival rates exceeded 60 % in both KO1 and KO2, compared with approximately 40 % in WT plants (Fig. 2F and G). Among the two alleles, KO1 exhibited the most pronounced osmotic stress tolerance phenotype, maintaining substantially more green tissue and reduced wilting throughout the treatment. Consistent with this phenotype, electrolyte leakage tended to be lower in both KO1 and KO2 than in WT plants at 2 d after PEG treatment (DT), although these differences did not reach statistical significance, and leakage levels became largely comparable among all genotypes by 4 DT. Although not statistically significant, this trend is consistent with the possibility that loss of *OsPSY8* may modestly slow the progression of cellular damage during osmotic stress (Fig. 2D). Together, these results demonstrate that OsPSY8 acts as a negative regulator of osmotic stress tolerance in rice.

### OsPSY8 promotes root growth under non-stress conditions

Previous studies showed that overexpression of *OsPSY1*, *OsPSY2*, and *OsPSY5* enhances root growth in rice (12, 13). To determine whether OsPSY8 similarly promotes root growth, we measured root length in *OsPSY8*-OX plants. OsPSY8-OX plants exhibited longer roots than WT plants (Fig. S3A and S3B). Although we hypothesized that loss of *OsPSY8* would result in shorter roots, *ospsy8* mutants did not exhibit shorter roots than WT plants (Fig. S3A and S3B). This observation suggested that functional redundancy within the *OsPSY* family might compensate for the loss of *OsPSY8*. Because functionally redundant genes often display similar expression patterns and compensatory upregulation following gene disruption (18), we examined their expression patterns and transcriptional responses to identify potential functional homologs of *OsPSY8*. A phylogenetic analysis based on promoter sequences revealed that *OsPSY5* and *OsPSY8* share closely related promoter regions (Fig. S3E), consistent with their similar spatial expression patterns and transcriptional responses to osmotic stress (Fig. 1A and 1C). Furthermore, expression analysis of all eight *OsPSY* genes in *ospsy8* mutants revealed a nearly twofold increase in *OsPSY5* expression, whereas the remaining family members were unchanged (Fig. S3F), suggesting compensatory upregulation of *OsPSY5* in the absence of *OsPSY8*. To directly test for functional redundancy between *OsPSY5* and *OsPSY8*, we generated an *ospsy5* knockout line (Fig. S3E) and crossed it with the *ospsy8* KO2 allele. Homozygous *ospsy5 ospsy8* double mutants were identified in the F2 generation, and F3 seeds were harvested for phenotypic analysis. Loss of *OsPSY5* alone did not significantly affect root length, even though previous work showed that loss of *OsPSY5* reduces root growth (13). The *ospsy8* displayed an increase in root length, similar to that observed for KO1 (Fig. S3A, B). Unexpectedly, the *ospsy5 ospsy8* double mutant also exhibited longer roots than WT (Fig. S3F, G). Notably, while WT and single-mutant seedlings developed crown roots by 5 days after germination, double-mutant seedlings instead developed a single elongated primary root, with apparently reduced crown root formation. The absence of a short-root phenotype in the *ospsy5 ospsy8* double mutant may reflect additional functional compensation by other root-expressed *OsPSY* family members, such as *OsPSY4* and *OsPSY6* (Fig. 1C), while the impaired crown root development suggests that the *OsPSY* family might regulate multiple components of root system architecture beyond root elongation. Collectively, these findings indicate that OsPSY8 contributes to root development under non-stress conditions, but its loss does not impair root elongation, likely due to broader functional redundancy within the OsPSY family.

### OsWRKY24 directly binds to and represses the OsPSY8 promoter

Constitutive *OsPSY8* expression promotes root growth but increases susceptibility to osmotic stress, whereas loss of *OsPSY8* enhances osmotic stress tolerance (Fig. S3 and 2), indicating that reduced OsPSY8 activity favors stress adaptation over growth. Given that *OsPSY8* expression is repressed under osmotic stress (Fig. 1C), we hypothesized that transcriptional repression of *OsPSY8* contributes to the transition from growth to stress adaptation. We therefore sought to identify transcription factors that regulate *OsPSY8* expression under osmotic stress conditions.

To identify transcription factors regulating *OsPSY8* expression, we first searched the RiceENCODE database for open chromatin regions within the *OsPSY8* promoter (19). Analysis of root Assay for Transposase-Accessible Chromatin with high-throughput sequencing (ATAC-seq) datasets identified three open chromatin regions within the 2.5-kb region upstream of the transcription start site (Fig. 3A). These regions were subsequently analyzed using PlantRegMap, which predicts TF-target interactions by scanning TF binding motifs in promoter sequences (20), and PlantPAN 4.0, which identifies TF binding sites within query sequences (21). To further narrow the candidate list, we used the plant single-cell database, scPlantDB (22), to identify TFs co-expressed with *OsPSY8*. scPlantDB detected *OsPSY8* expression in sclerenchyma cells, root epidermis, and root stele. Because lateral roots, where *OsPSY8* promoter activity was enriched in our GUS analysis (Fig. 1B), are not annotated as a separate cell-type category in this database, we used root stele expression as a biologically relevant proxy for lateral-root-associated expression, given that lateral root primordia are thought to initiate from cell layers within the stele (23). We then compared the root stele-enriched gene set with the candidate TF lists generated by PlantRegMap and PlantPAN 4.0 and found that OsWRKY24 was the only TF that satisfied all three criteria: predicted binding to the *OsPSY8* promoter by both databases and enriched expression in root stele cells.

**Figure 3.**
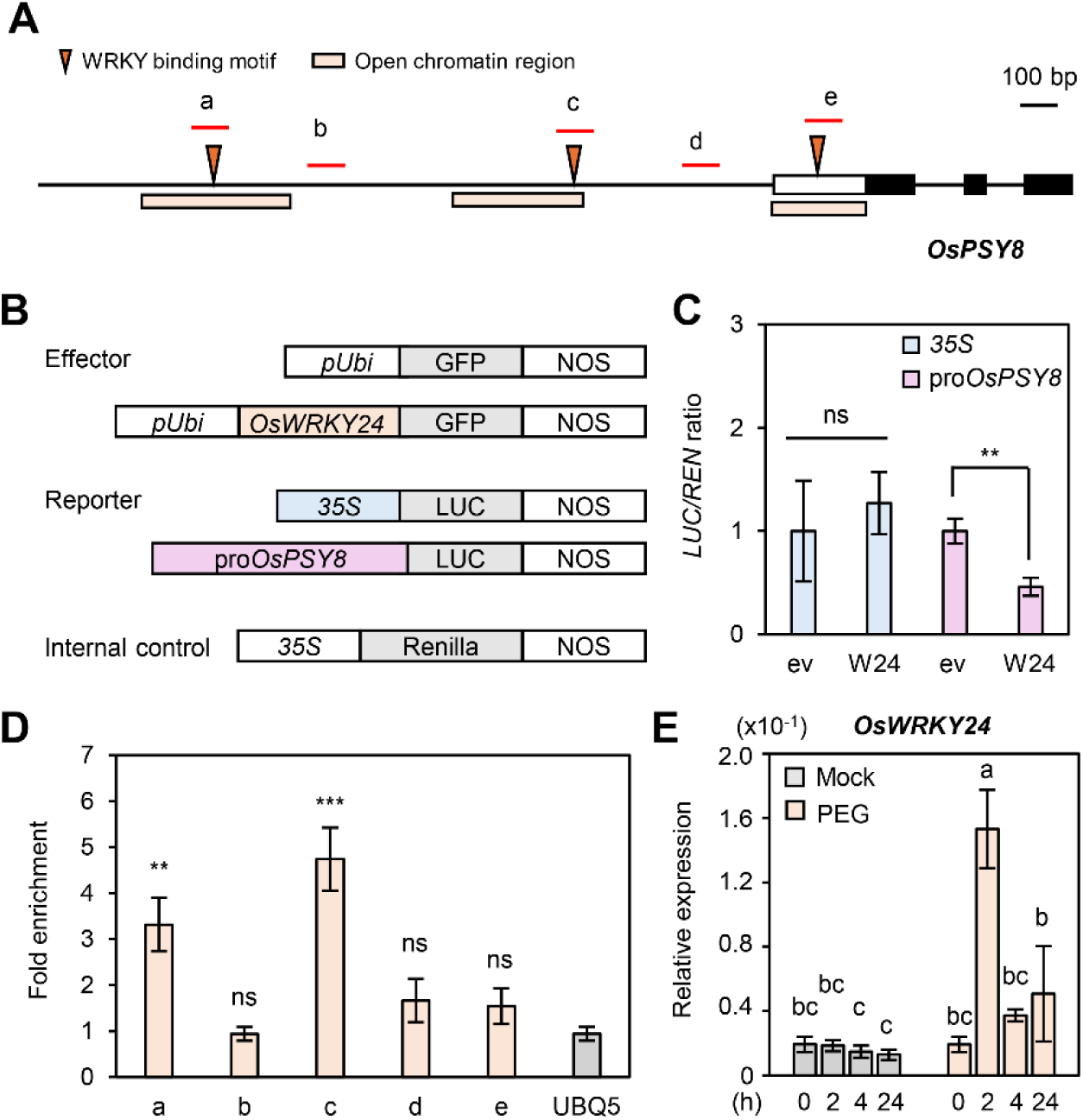
OsWRKY24 directly binds to and represses the *OsPSY8* promoter. (A) Schematic diagram of the *OsPSY8* genomic region showing the promoter region (line upstream of the 5′ UTR), 5′ UTR (open box), exons (filled black boxes), introns (lines connecting exons), the locations of open chromatin regions (orange boxes), WRKY binding motifs (W-box; orange arrowheads), and amplicons (a-e, red lines) used for ChIP-qPCR analysis. Scale bar, 100 bp. (B) Constructs used for the dual-luciferase (LUC) assay. Effector constructs consisted of *pUbi*::GFP (empty vector control) or *pUbi*::*OsWRKY24*-GFP. Reporter constructs consisted of *35S*::LUC or *pOsPSY8*::LUC. A *35S*::Renilla construct was used as an internal control. (C) LUC/REN ratio from dual-luciferase assays in rice protoplasts co-transfected with effector (empty vector (ev) or OsWRKY24 (W24)) and reporter (*35S*::LUC or *pOsPSY8*::LUC) constructs. (D) ChIP-qPCR analysis showing enrichment of OsWRKY24 at *OsPSY8* promoter amplicons a-e. The *OsUBQ5* gene was used as the negative control. Fold enrichment was calculated using the 2^−ΔΔCt^ method, where ΔCt = Ct(IP) − Ct(input), and ΔΔCt = ΔCt(OsWRKY24-GFP) − ΔCt(ev control). (C, D) Asterisks indicate statistically significant differences from control (\*\**P* < 0.01; two-tailed Student’s t-test). ns, not significant. (E) Expression levels of *OsWRKY24* in roots of WT plants treated with mock or PEG at 0, 2, 4, and 24 h. Relative expression was quantified by RT-qPCR as the transcript level of each target gene normalized to rice *OsUBQ5* using the 2^−ΔΔCT^ method. Data represent mean ± SD (n = 3 biological replicates), each replicate consisting of three plants. Statistical analyses were conducted using one-way ANOVA followed by Tukey’s honestly significant difference (HSD) post hoc test for multiple comparisons, with different letters indicating significant differences among groups (*P* < 0.05).

We first examined the effect of OsWRKY24 on *OsPSY8* promoter activity using a dual-luciferase (LUC) assay. A reporter construct containing the 2.5-kb promoter region upstream of the OsPSY8 transcription start site fused to LUC was cotransfected into rice protoplasts together with an OsWRKY24 expression construct (Fig. 3B). Cotransfection of *pUbi*::*OsWRKY24*-GFP significantly reduced LUC activity compared with the GFP control, indicating that OsWRKY24 represses *OsPSY8* promoter activity (Fig. 3C). To determine whether OsWRKY24 directly regulates *OsPSY8*, we performed chromatin immunoprecipitation (ChIP) assays. OsWRKY24 was significantly enriched at promoter amplicons a and c, which contain the WRKY-binding W-box motif (5′-TTGAC(C/T)-3′), demonstrating direct binding of OsWRKY24 to the *OsPSY8* promoter *in planta* (Fig. 3D). We next examined *OsWRKY24* expression in response to osmotic stress. *OsWRKY24* transcript levels in roots increased rapidly within 2 h of osmotic stress treatment, suggesting that the osmotic stress-induced decline in *OsPSY8* expression is preceded by increased *OsWRKY24* expression and direct repression of the *OsPSY8* promoter. (Fig. 3E). Together, these results identify OsWRKY24 as a direct transcriptional repressor of *OsPSY8* under osmotic stress.

### OsPSY8 modulates growth- and stress-associated transcriptional programs under osmotic stress

To explore the downstream signaling pathways regulated by OsPSY8 under osmotic stress, we compared transcriptome profiles among WT, OX (OX2), and KO (KO1) roots at 0 and 4 h after osmotic stress treatment. Sample reproducibility and separation were confirmed by principal component analysis (PCA) (Fig. S4A). Volcano plots of osmotic stress-treated WT revealed a large-scale transcriptional response, with 3,607 and 3,108 genes differentially expressed at 0 h and 4 h, respectively, using a threshold of |log₂FC| > 1 and a false discovery rate (FDR)-adjusted *P* ≤ 0.05 (Fig. S4B and Dataset S1). Against this broad transcriptomic background, loss or overexpression of *OsPSY8* induced a comparatively specific set of transcriptional changes: KO altered 139 and 335 genes at 0 h and 4 h (Fig. S4C, D, and Dataset S2, S3), and OX affected 90 and 191 genes at the respective time points (Fig. S4E, F and Dataset S4, S5).

Functional classification revealed distinct transcriptional programs consistent with the phenotypes of KO and OX plants. KO plants showed upregulation of stress-responsive genes associated with osmotic stress adaptation, including genes involved in lignin biosynthesis, ROS scavenging, and compatible solute biosynthesis (Fig. 4A). Expression of *OsCCR15* (*Cinnamoyl-CoA Reductase 15*), a member of the rice *CCR* gene family implicated in the monolignol pathway for lignin biosynthesis (24), was validated by qRT-PCR and was elevated in both KO alleles (Fig. 4C). We additionally determined whether the altered gene expression observed in KO roots could be rescued by exogenous OsPSY8 treatment. Before treatment, the bioactivity of the synthetic OsPSY8 peptide (see SI Appendix, Materials and Methods for peptide sequence, sulfation site, and synthesis details) was first confirmed in *Arabidopsis tyrosylprotein sulfotransferase* (*tpst*) mutant, which lacks the TPST required for tyrosine sulfation and therefore lacks biologically active PSY peptides and typically exhibits shorter primary roots than Col-0 (11,17). Treatment with 500 nM OsPSY8 increased root length of *tpst* mutants (Fig. S5A and B). Next, KO1 and KO2 roots were treated with 500 nM, and the expression of *OsCCR15* and OsGSTU36 (*Tau-class Glutathione S-Transferase 36*, involved in cellular detoxification processes, (25)) was measured by qRT-PCR. After 2 h of peptide treatment, *OsCCR15* and *OsGSTU36* expression showed a trend toward reduction in KO roots relative to mock-treated KO roots, although most reductions were not statistically significant and expression was not fully restored to mock-treated WT levels (Fig. S5C and D). This partial response to exogenous OsPSY8 peptide in the KO plants provides indirect evidence that the loss of endogenous bioactive OsPSY8 may contribute to the increased expression of stress-responsive genes in KO plants.

**Figure 4.**
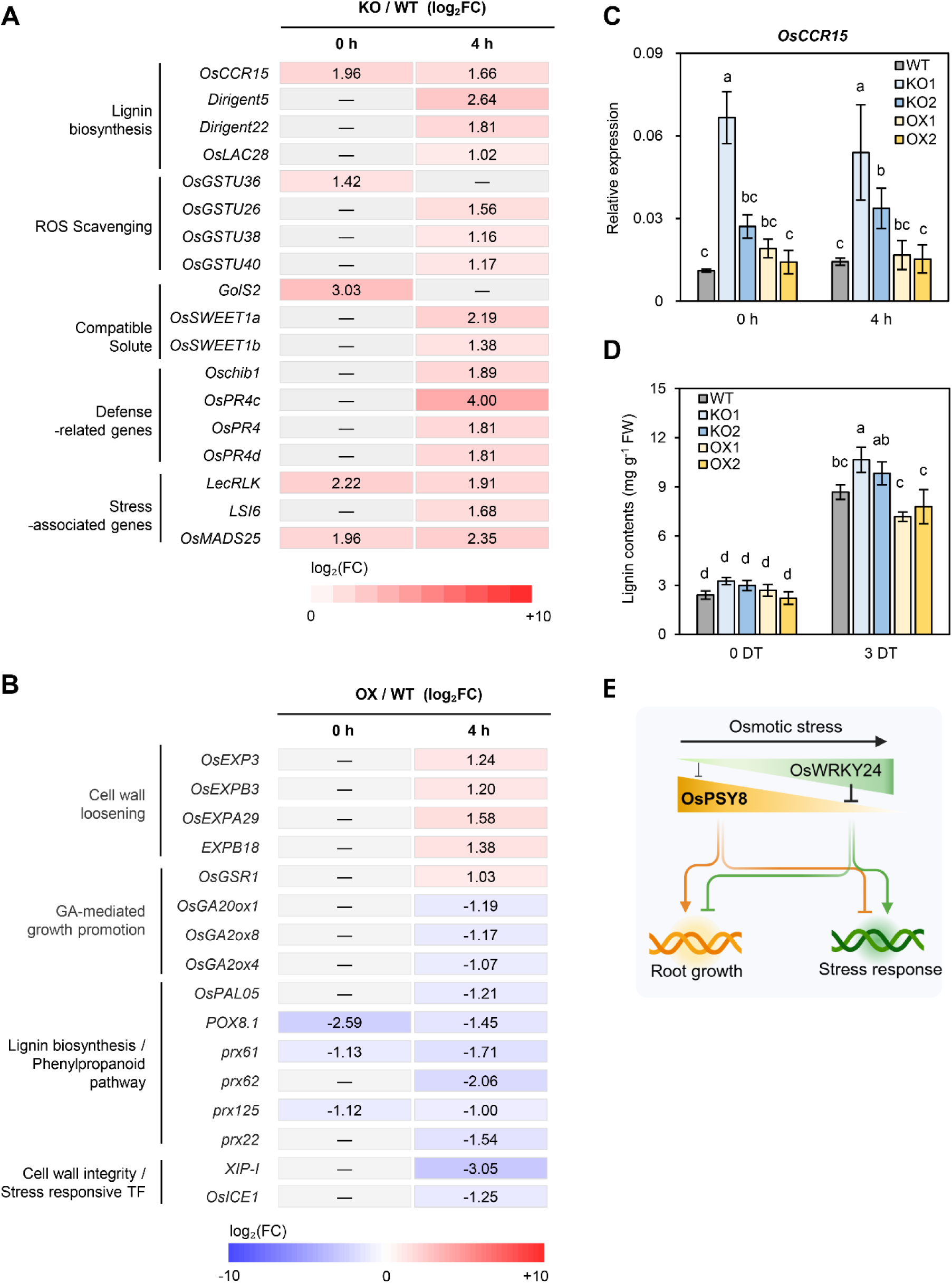
OsPSY8 modulates growth- and stress-associated transcriptional programs under osmotic stress. (A) Heatmap showing log₂fold changes (log₂FC) of selected stress-responsive genes upregulated in KO relative to WT at 0 h and 4 h after osmotic stress treatment. Genes are grouped by functional categories. Dashes indicate non-significant changes. (B) Heatmap showing log₂FC of selected growth-associated and stress-responsive genes differentially expressed in OX relative to WT at 0 h and 4 h after osmotic stress treatment. Genes are grouped by functional category. Dashes indicate non-significant changes. (C) Expression level of *OsCCR15* in roots of WT, KO1, KO2, OX1, and OX2 plants at 0 h and 4 h after osmotic stress treatment. Relative expression was quantified by RT-qPCR as the transcript level of each target gene normalized to rice *OsUBQ5* using the 2^−ΔΔCT^ method. Data are mean ± SD (n = 3 biological replicates), each replicate consisting of three plants. (D) Lignin contents in roots of WT, KO1, KO2, OX1, and OX2 plants at 0 and 3 days after osmotic stress treatment (DT). Data are mean ± SD (n = 3). (C, D) Statistical analyses were conducted using one-way ANOVA followed by Tukey’s honestly significant difference (HSD) post hoc test for multiple comparisons, with different letters indicating significant differences among groups (*P* < 0.05). (E) Working model for OsPSY8 function. OsPSY8 promotes root growth while suppressing stress-responsive pathways under non-stress conditions. Osmotic stress induces OsWRKY24, which represses *OsPSY8* expression, thereby suppressing growth-associated transcriptional programs and activating stress-responsive programs.

OX plants showed reduced expression of a subset of lignin biosynthesis genes before stress, and this repression became more extensive after osmotic stress treatment, including reduced expression of *OsPAL05* gene encoding phenylalanine ammonia-lyases, which initiate the phenylpropanoid pathway (Fig. 4B). Under osmotic stress, OX plants maintained a growth-promoting transcriptional program, with sustained expression of cell wall-loosening expansins and reduced expression of gibberellin (GA)-catabolic *OsGAox* genes, a pattern consistent with reduced GA catabolism and continued growth under stress (Fig. 4B). To determine whether these transcriptional changes were accompanied by altered lignin accumulation, we quantified lignin content in roots before and after osmotic stress. Before stress treatment, lignin content did not differ significantly among genotypes, despite transcriptional changes in lignin biosynthesis genes (Fig. 4D). After stress treatment, lignin content tended to be higher in KO plants than in WT, with a significant increase detected only in KO1 (Fig. 4D). In contrast, OX plants showed a trend toward lower lignin accumulation than WT after osmotic stress, although these differences were not statistically significant (Fig. 4D). Of note, *OsPSY1*-overexpressing plants were also reported to have reduced lignin accumulation in roots (12), suggesting that altered PSY signaling may influence root lignin deposition. Taken together, these results indicate that constitutive expression of *OsPSY8* suppresses stress-responsive pathways while promoting growth-associated transcriptional programs, whereas loss of *OsPSY8* function activates stress-responsive pathways (Fig. 4E).

## Discussion

Rice is a staple food for more than half of the world’s population, and the rising frequency and severity of drought events increasingly threaten its production. Understanding the molecular mechanisms that enable rice plants to transition from growth to stress adaptation is therefore essential for developing climate-resilient crops. In this study, we demonstrate that OsPSY8 functions as a regulator of the growth-to-stress transition in rice roots. *OsPSY8* is predominantly expressed in root tissues, where it promotes root growth and suppresses stress responses under non-stress conditions. Upon osmotic stress, OsWRKY24 expression is upregulated, and OsWRKY24 binds the *OsPSY8* promoter to repress *OsPSY8* expression. This mechanism shifts the plant to stress response mode rather than growth (Fig. 3C-E). Consistent with this shift, *ospsy8* mutants showed increased expression of lignin biosynthesis genes, accompanied by a trend toward greater lignin accumulation after osmotic stress, although a statistical significance was detected only in KO1. Such changes in lignin deposition could potentially contribute to strengthening root cell walls and apoplastic barriers during osmotic stress. These findings expand our understanding of PSY peptide signaling in rice and demonstrate that osmotic stress-induced repression of *OsPSY8* contributes to the transition from growth to stress adaptation.

Previous studies in *Arabidopsis* showed that exogenous application of the PSY5 peptide promotes root elongation, and loss of its cognate receptors phenocopies this response, demonstrating that PSY peptides promote growth by repressing PSYR signaling (14). In addition, either treatment of PSY peptide or loss of PSYR function compromises plant tolerance to abiotic stresses, suggesting that a constitutive PSY response represses PSYR-mediated stress responses. This ligand-deprivation-dependent activation of stress responses is a distinctive feature of PSY signaling, because most RLKs activate receptor signaling upon ligand binding (26). Our findings reveal a similar functional pattern in rice, in which constitutive *OsPSY8* expression maintains growth-associated programs while attenuating stress responses, whereas repression of *OsPSY8* under osmotic stress induces stress-responsive programs (Fig. 4E). Interestingly, in an amino acid sequence-based phylogenetic analysis of PSY proteins from multiple plant species (33), OsPSY8 appeared most closely related to PSY5, which was used to demonstrate ligand-deprivation-dependent activation of stress responses in *Arabidopsis* (14). Currently, it remains unclear whether a similar PSY signaling mechanism also operates in rice, as described in Arabidopsis. We recently proposed two putative rice PSYR-like receptors, *OsPSYR1* (LOC_Os07g05740) and *OsPSYR2* (LOC_Os04g42700), based on homology to *Arabidopsis* PSYRs (27). Notably, *OsPSYR2* is predominantly expressed in root tissue, coinciding with the root-predominant expression of *OsPSY8* (Fig. S6). However, functional analysis of these candidates, such as direct receptor-ligand interaction and characterization of *ospsyr* mutants will be required to determine whether ligand-deprivation-dependent activation of stress responses is conserved in rice.The eight OsPSY genes exhibited distinct transcriptional responses to osmotic stress; *OsPSY5* and OsPSY8 were downregulated, whereas the remaining six family members (*OsPSY1*, *OsPSY2*, *OsPSY3*, *OsPSY4*, *OsPSY6*, and *OsPSY7*) were induced (Fig. 1C). These observations suggest that osmotic stress alters the overall composition of the PSY peptide pool rather than simply reducing OsPSY8 abundance. This stress-responsive expression pattern contrasts with that of *Arabidopsis PSY* genes, which are not stress inducible and maintain largely stable expression under various stress conditions (14). This difference suggests that rice regulates PSY signaling through stress-dependent transcriptional control of PSY peptide genes, unlike Arabidopsis. Notably, although six *OsPSY* genes were induced by osmotic stress, *OsPSY5* and *OsPSY8* are the predominant root-expressed family members under non-stress conditions (Fig. 1A), suggesting that repression of these abundant members may have a greater impact on the bioactive PSY peptide pool than induction of lower-abundance family members. Despite the higher basal expression of *OsPSY5* than *OsPSY8*, the alteration of *OsPSY8* expression alone affected osmotic stress tolerance, suggesting a specialized role for OsPSY8 in osmotic stress responses. Such functional differences may reflect their distinct cell-type-specific expression patterns or receptor interactions, which remain to be determined. Furthermore, defining the distinct functions of OsPSY5 and OsPSY8 will require characterization of plants expressing each gene under root-specific promoters, thereby minimizing potential ectopic effects associated with the constitutive promoter used in this study.

*OsPSY8* is directly repressed by OsWRKY24, a transcription factor previously shown to positively regulate resistance against *Magnaporthe grisea*, the causal agent of rice blast disease (28). Although the role of OsWRKY24 has been studied in foliar disease resistance in rice, *WRKY25*, *WRKY26*, and *WRKY33*, the Arabidopsis orthologs of *OsWRKY24* identified using the RAP-DB ortholog search tool based on OrthoFinder orthogroup clustering, have been implicated in abiotic stress tolerance. The *wrky33* loss-of-function mutant exhibits increased sensitivity to drought (29), whereas *WRKY25*, *WRKY26*, and *WRKY33* act partially redundantly to positively regulate thermotolerance, with loss-of-function mutants showing increased heat stress sensitivity (30). Future analysis of *OsWRKY24* overexpression and loss-of-function lines under osmotic stress will be important to determine whether OsWRKY24 activity is required for OsPSY8-dependent stress responses. ChIP-qPCR in OsWRKY24-tagged plants exposed to osmotic stress would further test whether OsWRKY24 binding to the *OsPSY8* promoter occurs specifically under stress conditions *in planta*.

A recurring challenge in engineering stress-tolerant crops is that constitutive activation of stress responses often compromises plant growth because of the associated energy costs (2, 31). In contrast, disruption of *OsPSY8* enhanced osmotic stress tolerance without detectable defects in root growth, possibly because of functional compensation by the closely related gene *OsPSY5* (Fig S3). The diverse expression patterns of the eight rice *PSY* genes further suggest a combination of functional specialization and redundancy within the family. A better understanding of these relationships may enable more precise manipulation of PSY signaling pathways to enhance stress resilience while minimizing growth penalties.

In addition to drought tolerance, PSY peptides represent attractive targets for engineering because they have the potential to influence plant-microbiome interactions (32). Overexpression of *OsPSY1* has been reported to alter root exudate composition and reduce methane emissions from rice paddies (12), indicating that PSY-family peptides can modulate the chemical interface between roots and the rhizosphere. Recent studies in sorghum have shown that drought-induced changes in root metabolite and exudate composition are closely associated with shifts in root-associated microbial communities, suggesting that drought-responsive exudates contribute to microbiome restructuring under water deficit conditions (33). Future studies examining how altered OsPSY8 signaling affects root exudate composition and microbial community assembly under drought conditions may provide additional insights into the broader ecological functions of PSY peptides.

In this study, we used PEG-induced osmotic stress to investigate the role of OsPSY8 in coordinating growth and stress responses, providing a biological foundation for future validation under soil-drying conditions. To translate these mechanistic insights into crop improvement strategies, future studies should investigate how modulation of OsPSY8 signaling affects plant performance under soil-grown conditions, including greenhouse and field environments. Such investigations will be essential to determine whether engineering the OsPSY8 pathway can improve drought resilience while maintaining crop productivity under agronomically relevant conditions. Furthermore, given that PSY peptides are conserved across diverse crop species, including maize, soybean, and wheat (34, 35), a key next step will be to determine whether PSY-mediated regulation of the growth-to-stress transition operates similarly in other major crops.

## Materials and Methods

### Plant MaterialsGrowth Conditions, and Stress Treatment

*Oryza sativa* ssp*. Japonica* cultivar Kitaake was used as the WT background throughout this study. Seeds were germinated in water at 28 °C for 3 d and subsequently grown hydroponically in Hoagland solution (36). Seedlings of different genotypes were grown in separate hydroponic containers throughout the experiment. Seedlings were grown in a growth chamber under a 14-h light/10-h dark photoperiod at 26 °C/24 °C (day/night).

For osmotic stress survival assays, 9-day-old hydroponically grown seedlings were transferred to Hoagland solution containing 25 % (w/v) PEG6000. Seedlings were treated for 4 d in the experiment shown in Fig. 2 and for 5 d in the experiment shown in Fig. S2. The PEG solution was then replaced with fresh Hoagland solution, and survival rates were measured after 7 d of recovery.

### Histochemical GUS staining

The promoter regions of *OsPSY5* and *OsPSY8* were amplified using primers described in Table S1 and inserted upstream of the GUS gene in pMDC163, a Gateway binary vector. The plasmid was transformed into *Agrobacterium* strain LBA4404, and transgenic plants were generated following a previously described method (37). GUS staining was performed as previously described (38). The 9-day-old transgenic plants were incubated in GUS staining solution (50 mM phosphate buffer, 1 mM potassium ferricyanide, 1 mM potassium ferrocyanide, 2 mM X-Gluc, 0.1% Triton X-100, 10 mM EDTA [pH 8.0], and MilliQ water) at 37 °C for 48 h. The samples were observed under a stereomicroscope (ZEISS SteREO Discovery.V20). The root tissues were divided into 20 mm segments and embedded in 4 % low-melting-point agarose. Horizontal sections were obtained using a Vibratome VT 1200S (Leica) and imaged with a stereomicroscope.

### RT-qPCR Analysis

Total RNA was extracted from rice tissues using TRIzol reagent (Invitrogen) and treated with the TURBO DNA-free Kit (Ambion) to remove residual genomic DNA. First-strand cDNA was synthesized using the High-Capacity cDNA Reverse Transcription Kit (Applied Biosystems). Quantitative PCR was performed on a CFX96 Real-Time PCR System coupled with a C1000 Thermal Cycler (Bio-Rad) using iTaq Universal SYBR Green Supermix (Bio-Rad). Relative transcript levels were normalized to *OsUBQ5*. The number of biological replicates analyzed for each experiment is indicated in the corresponding figure legend Primers used in this study are listed in Table S1.

### Generation of Transgenic Rice Lines

For the generation of overexpression lines, the genomic *OsPSY5* and *OsPSY8* sequence was amplified using the primers listed in Table S1 and cloned into the pC1300-Ubi-Nos vector, in which transgene expression is driven by the *UBIQUITIN* (*Ubi*) promoter. Plasmid construction and rice transformation were performed as previously described (12, 37). For the generation of the CRISPR/Cas9-mediated knockout mutants, custom-designed guide RNAs (gRNAs) (Table S1) targeting *OsPSY8* coding regions were cloned into the pOs-sgRNA vector and subsequently introduced into the destination vector pH-Ubi-Cas9-7 through Gateway recombination (39). Transgenic plants were generated following a previously described method (37) using *Agrobacterium* strain EHA105 carrying the binary vector. All transgenic lines used in this study were confirmed as single-copy homozygous insertions by hygromycin segregation analysis and were advanced to at least the T2 generation before phenotypic analysis.

### Electrolyte leakage assay

Fully expanded leaves from 9-day-old WT, *OsPSY8*-OX, and *ospsy8* mutant seedlings were sampled before, 2, and 4 days after PEG treatment. Samples were thoroughly rinsed with Milli-Q water and incubated in 8 mL Milli-Q water for 3 h with gentle inversion, after which initial conductivity was measured using a YSI 3200 conductivity instrument. Samples were then boiled at 85 °C for 20 min, cooled to room temperature, and total conductivity was measured. Electrolyte leakage rate (%) was calculated as (initial conductivity / total conductivity) × 100. Data represent mean ± SD of four biological replicates.

### Dual-Luciferase Assay in Rice Protoplasts

Dual-luciferase (LUC) assays were performed in rice protoplasts as previously described (40). A 2-kb promoter fragment of *OsPSY8* was cloned into the pJD301 vector (41) using primers in Table S1 upstream of the LUC reporter gene, and *pUbi*::*OsWRKY24*-GFP was used as the effector construct. pEGB *35S*::Renilla:Tnos (GB0109; Addgene) was used as an internal control vector. Reporter, effector, and control plasmids were cotransfected into rice protoplasts by PEG-mediated transfection. After incubation in the dark for 14 h, firefly and Renilla luciferase activities were measured using a Dual-Luciferase Reporter Assay System (Promega), and promoter activity was expressed as the ratio of firefly luciferase to Renilla luciferase activity.

### ChIP assay

ChIP assays were performed as previously described (40). Rice protoplasts expressing OsWRKY24-GFP or GFP alone were crosslinked, and chromatin was isolated, sonicated, and immunoprecipitated using an anti-GFP antibody (Santa Cruz Biotechnology). Binding of OsWRKY24 to the *OsPSY8* promoter was quantified by qPCR using immunoprecipitated DNA. *OsUBQ5* served as a negative control.

### RNA-seq analysis

Total RNA was extracted using the GeneJET Plant RNA Purification Kit (Thermo Fisher Scientific) according to the manufacturer’s instructions. RNA sequencing was performed using three biological replicates from each genotype and treatment condition, including WT, OX2, and KO1 roots sampled before and 4 h after PEG treatment. RNA-seq reads were quality-filtered using fastp v0.24.0 and aligned to the reference genome using STAR v2.7.11, with unique molecular identifier (UMI)-based deduplication performed using UMIcollapse v1.1.0. Gene-level counts were quantified using featureCounts (subread v2.1.1), and differential expression analysis was performed using edgeR v4.0.16.

### Lignin quantification

Lignin content was quantified using a modified thioglycolic acid assay as previously described (42). Approximately 100 mg of fresh root tissue was ground in liquid nitrogen, extracted with methanol, and incubated with thioglycolic acid and HCl at 80 °C to generate lignin–thioglycolic acid complexes. The complexes were extracted with 1 M NaOH, precipitated by acidification, and subsequently resuspended in 1 M NaOH. Absorbance was measured at 280 nm, and lignin content was calculated using an alkali lignin (Sigma-Aldrich, CAS No. 8068-05-1) standard curve.

### Use of Artificial Intelligence

ChatGPT (OpenAI) was used solely for language editing and improvement of manuscript phrasing. All scientific content, analyses, interpretations, and conclusions were developed and verified by the authors.

## Author Contributions

Y.S. designed the research, performed experiments, analyzed the data, and wrote the manuscript; E.Y.R. revised the manuscript; J.C.-Y.L. assisted with GUS staining experiments and revised the manuscript; M.C. and G.A. generated *ospsy5*, *ospsy8* knockout and GUS reporter transgenic plants; S.S.K. and P.W.C synthesized the sulfated OsPSY8 peptide under the supervision of R.J.P; M.F.E. generated rice lines overexpressing OsPSY5 and OsPSY8 and revised the manuscript; and P.C.R. supervised the study, provided funding and resources, and revised the manuscript.

## Classification

Major category: Biological Sciences

Minor category: Agricultural Sciences

## Acknowledgments

We thank Valley Stewart for valuable guidance and mentorship throughout this study. We also thank Sanyukta Rohom, Jiayi An, and Jinyoung Kwon for assistance with genotyping and phenotyping experiments. This work was supported by NIH (1R35GM148173) to P.C.R. and the Joint BioEnergy Institute. The work conducted at the Joint BioEnergy Institute was supported by the U.S. Department of Energy, Office of Science, Biological and Environmental Research Program, through contract DE-AC02-05CH11231 between Lawrence Berkeley National Laboratory and the U.S. Department of Energy. The United States Government retains and the publisher, by accepting the article for publication, acknowledges that the United States Government retains a nonexclusive, paid-up, irrevocable, worldwide license to publish or reproduce the published form of this manuscript, or allow others to do so, for United States Government purposes. Any subjective views or opinions that might be expressed in this paper do not necessarily represent the views of the U.S. Department of Energy or the United States Government.

