## Supplementary material for "Signaling by a tyrosine-sulfated peptide balances growth and osmotic stress response in rice": SI appendix

**This PDF file includes:**

Supporting text

Figures S1 to S6

Table S1

Legends for Datasets S1 to S6

SI References

**Other supporting materials for this manuscript include the following:**

Datasets S1 to S5

Supporting Information Text

**Materials and Methods**

**Sulfated peptide synthesis and treatment.**

Sulfated OsPSY8 peptides (DY(SO_3_)GQASANNRHNPHP) were prepared (50 µmol) on a SYRO I automated peptide synthesizer using standard Fmoc solid-phase peptide synthesis (SPPS) on 2-chlorotrityl chloride (2-CTC) resin (0.8-1.5 mmol/g, Novabiochem). The C-terminal amino acid was loaded on 2-CTC resin by treating the resin with a solution of Fmoc-Xaa-OH (3 equiv. relative to maximum loading) and iPr2NEt (8 equiv.) in CH_2_Cl_2_ (0.3 M concentration of Fmoc-Xaa-OH) for 16 h at rt. Following the loading of the C-terminal amino acid, Fmoc deprotection was performed with 40 % piperidine in DMF (3 mL, 3 min), followed by a second treatment with 20 % piperidine in DMF (3 mL, 10 min) at room temperature, and the resin was washed with DMF (× 4). Couplings were performed by treating the resin with a solution of Fmoc-Xaa-OH (4 equiv., 0.17 M), Oxyma (4.4 equiv., 0.18 M), and DIC (4 equiv., 0.17 M) in DMF at 40 °C. Sulfated tyrosine was introduced by coupling Fmoc-Tyr(SO₃nP)-OH (nP = neopentyl protecting group to avoid premature loss of sulfate during acidolytic cleavage) using standard DIC/Oxyma coupling conditions at room temperature. Fmoc-Tyr(SO₃nP)-OH was synthesized using the literature protocol (1). After DMF washes, a capping step was performed before the next SPPS cycle by treating the resin with acetic anhydride (5 vol.%) and iPr2NEt (10 vol.%) in DMF (3 mL) at room temperature, followed by DMF washes. Upon elongation, the peptide was subjected to acidolytic cleavage using TFA/*i*Pr_3_SiH/H₂O (90:5:5, v/v/v) for 2 h at room temperature. Upon concentrating the cleavage cocktail under vacuum, the crude peptide bearing a nP-protected sulfotyrosine was then precipitated using cold diethyl ether and collected by centrifugation (4,000 × g, 10 min, 4 °C). The nP deprotection was then performed by dissolving the peptide in a solution of 6 M Gdn•HCl, 1 M NH_4_OAc, and 0.1 M Na_2_HPO_4_ and incubating it for 1 h at 40 °C. After reverse-phase HPLC (RP-HPLC) purification and lyophilization, the sulfated peptide was isolated as a fluffy, white solid. Peptide OsPSY8 (20 µmol) was purified on a preparative C18 column (gradient: 0 to 50 % MeCN over 50 min, with 0.1 % NH4OH). Yield = 5.0 mg, 14 %.

For synthetic peptide treatments, sterilized seeds of the *Arabidopsis* *tpst* mutant (SALK_009847) were surface-sterilized with 70 % ethanol, stratified at 4°C for 2 days in the dark, and sown on MS medium (pH 5.7) supplemented with 0.3 % Gelzan (G024; Caisson Labs) and 500 nM sulfated OsPSY8 peptide, or an equal volume of water as a control. The peptide was diluted in Milli-Q water before addition to the medium. Seedlings were grown vertically under long-day conditions (16 h light/8 h dark) at 21 °C, and primary root length was measured at 4, 6, 8, 12, and 14 d after germination.

Figures

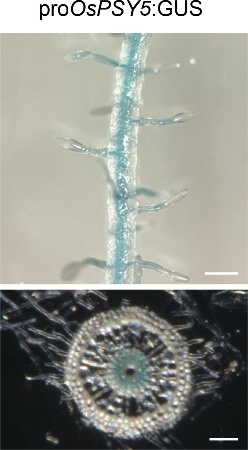

Fig. S1. Histochemical GUS staining of *pOsPSY5*::GUS transgenic rice seedlings. Samples were collected from 9-day-old seedlings. Upper panels, root differentiation zone images, scale bar = 500 µm; lower panels, 100-µm transverse sections prepared by vibratome sectioning, scale bar = 100 µm.

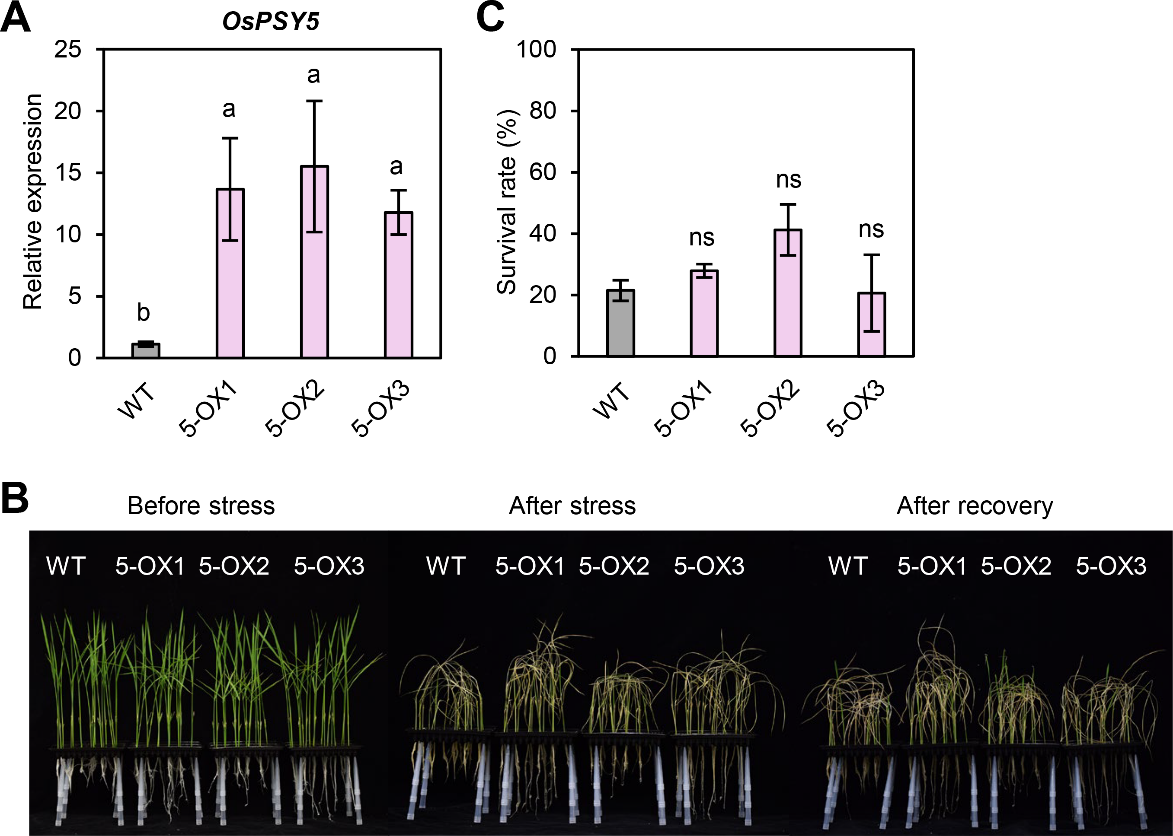

Fig. S2. *OsPSY5* overexpression does not confer drought stress tolerance in rice. (A) Relative expression levels of *OsPSY5* in WT and three independent *OsPSY5*-OX lines (5-OX1, 5-OX2, and 5-OX3~~#~~). Relative expression was quantified by RT-qPCR as the transcript level of each target gene normalized to rice *OsUBQ*) using the 2^-ΔΔCT^ method. Data represent mean ± SD (n = 4 biological replicates). Each replicate consists of three plants. Statistical analyses were performed using one-way ANOVA followed by Duncan's multiple range test, with different letters indicating significant differences among groups (*P* < 0.05). (B) Representative photographs of WT and *OsPSY5*-OX plants before stress, 5 days after PEG-induced osmotic stress, and 7 days after recovery. (C) Survival rates of WT and *OsPSY5*-OX lines (5-OX1, 5-OX2, and 5-OX3) after recovery. Each biological replicate consisted of one hydroponic box containing 40 plants. ns, not significant compared to WT (Student's *t*-test). Data are means ± SD (n = 3 biological replicates).

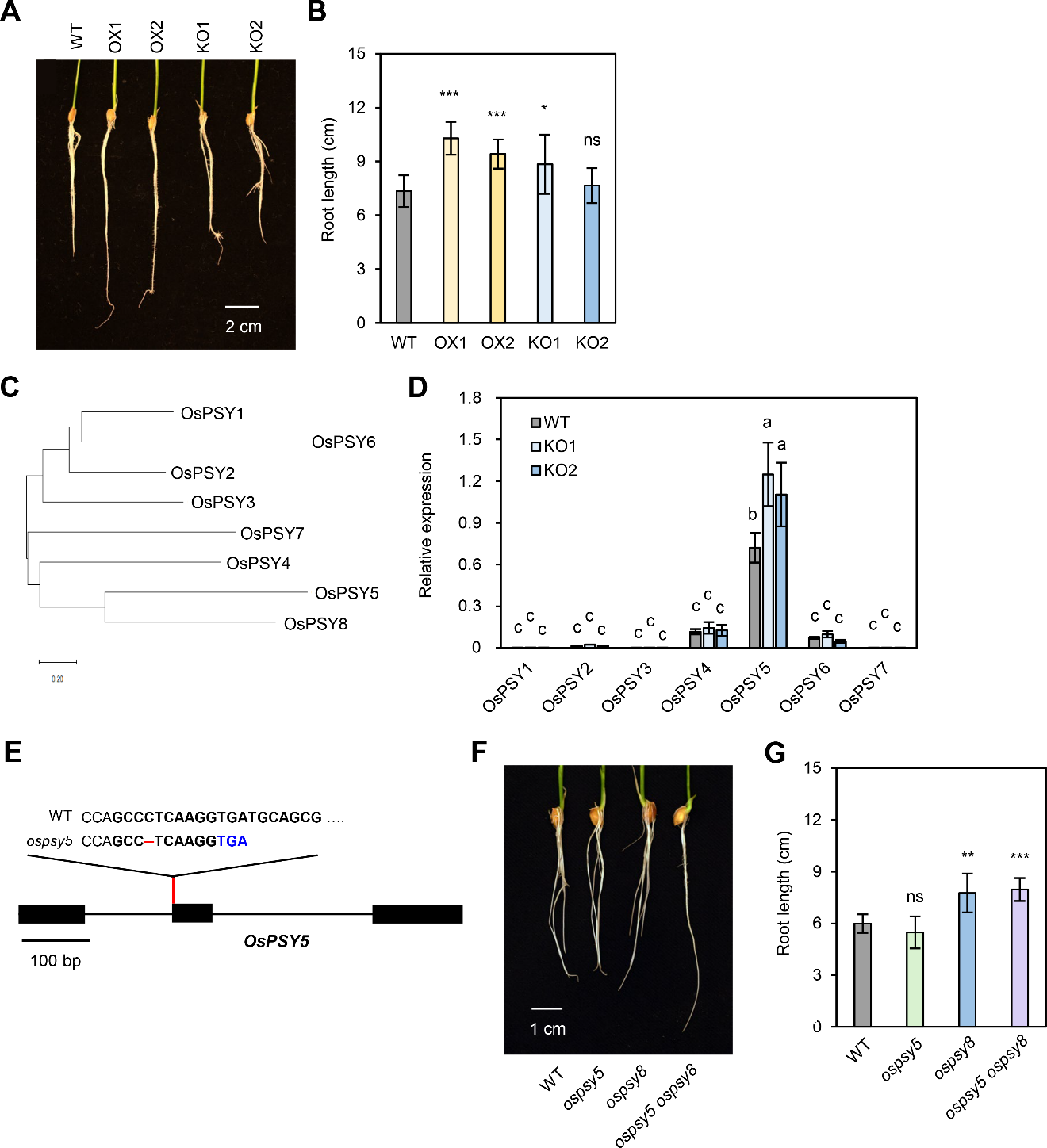

**Fig. S3. OsPSY8 promotes root growth under non-stress conditions.** (A) Representative images of roots from 9-day-old WT, OX, and KO plants. Scale bar, 2 cm. (B) Root length of WT, OX, and KO plants. Data are mean ± SD (n ≥ 10). (C) Phylogenetic tree of *OsPSY* family members inferred from promoter sequences (2.5-kb upstream from the transcription start site) using the Maximum Likelihood method with the General Time Reversible model. Branch lengths represent the number of substitutions per site. Evolutionary analyses were conducted in MEGA11 (2). (D) Relative expression levels of *OsPSY1*-*OsPSY7* in WT, KO1, and KO2 roots. Relative expression was quantified by RT-qPCR as the transcript level of each target gene normalized to rice *OsUBQ5* using the 2^-ΔΔCT^ method. Data are mean ± SD (n = 4 biological replicates), each replicate consisting of three plants. Statistical analyses were conducted using one-way ANOVA followed by Tukey's honestly significant difference (HSD) post hoc test for multiple comparisons, with different letters indicating significant differences among groups (*P* < 0.05). (E) Schematic diagram of the *OsPSY5* gene structure showing the 1-bp deletion in *ospsy5* that introduced a premature stop codon (TGA, blue). The red line indicates the deletion site. exons (filled black boxes), introns (lines connecting exons) Scale bar, 100 bp. (F) Representative images of roots from 5-day-old WT, *ospsy5*, *ospsy8* (KO2), and *ospsy5 ospsy8* double mutant. Scale bar, 1 cm.  (G) Root length of WT, *ospsy5*, *ospsy8* (KO2), and *ospsy5 ospsy8* double mutants. Data are mean ± SD (n = 8). (B, G) Asterisks indicate statistically significant differences from WT or Mock control (**P* < 0.05, ****P* < 0.001; two-tailed Student's *t*-test). ns, not significant.

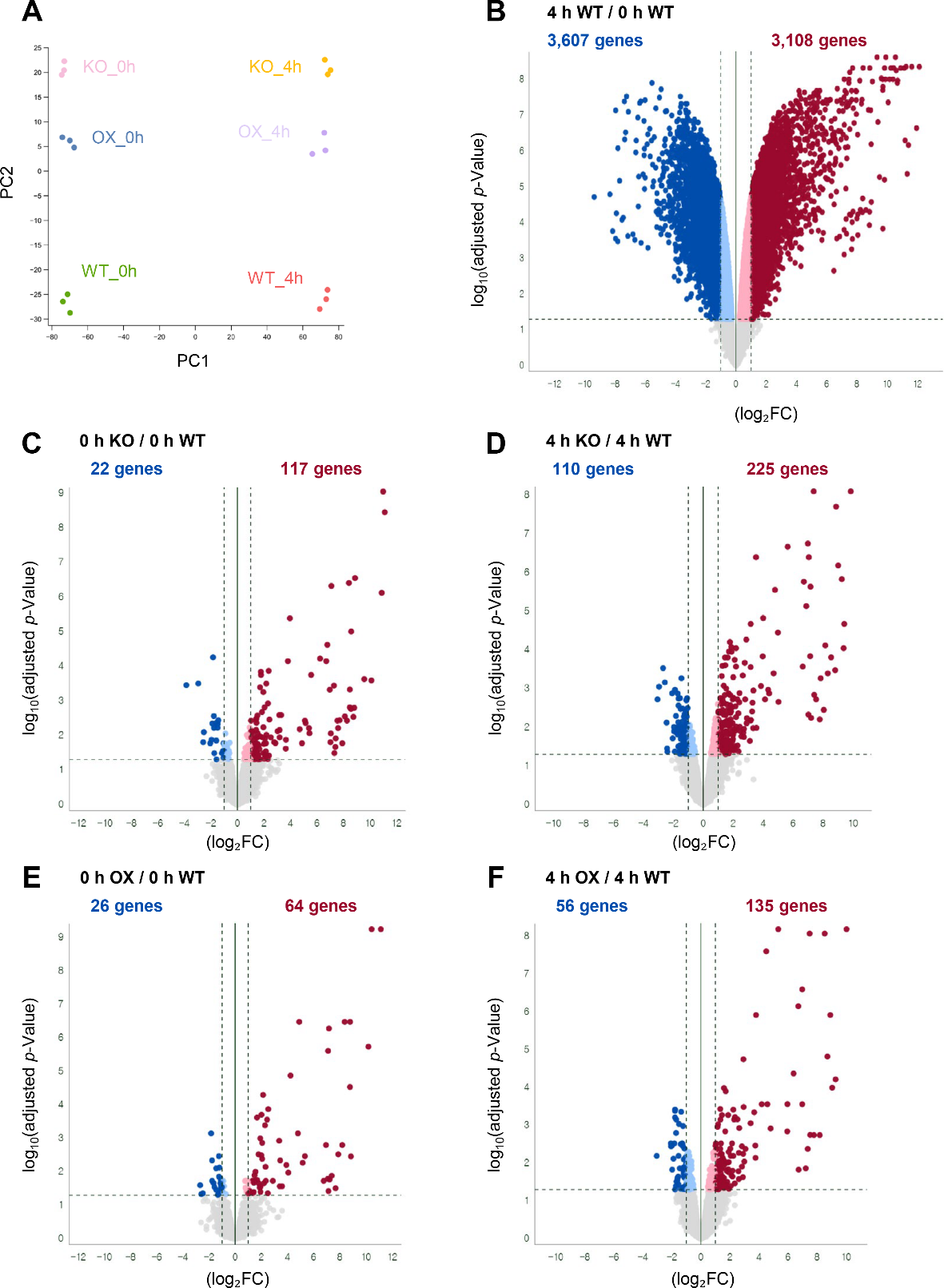

**Fig. S4. Principal component analysis (PCA) and volcano plots of RNA-seq data from WT, *ospsy8-1* (KO), and *OsPSY8*-OX #10 (OX) roots before and after osmotic stress.** (A) PCA was performed on the trimmed mean of M-values (TMM)-normalized counts with Pearson correlation from 18 samples representing three genotypes (WT, KO, and OX) under 0 h and 4 h after osmotic stress, with three biological replicates per group. PC1 (84.7% of variance explained) separates samples by treatment condition, and PC2 (6.3% of variance explained) further resolves the three genotypes within each treatment group. (B) Volcano plot comparing transcriptome profiles of WT roots at 0 h and 4 h after osmotic stress. (C, D) Volcano plots comparing transcript levels between WT and KO roots at (C) 0 h and (D) 4 h after osmotic stress. (E, F) Volcano plots comparing transcript levels between WT and OX roots at (E) 0 h and (F) 4 h after osmotic stress. The x- and y-axes represent log₂FC and log_10_(*P*-value), respectively. The blue and red dots represent downregulated DEGs with log₂(FC) < −1 and upregulated DEGs with log₂(FC) >1, respectively. Gray dots represent no significant differences in the transcriptomes.

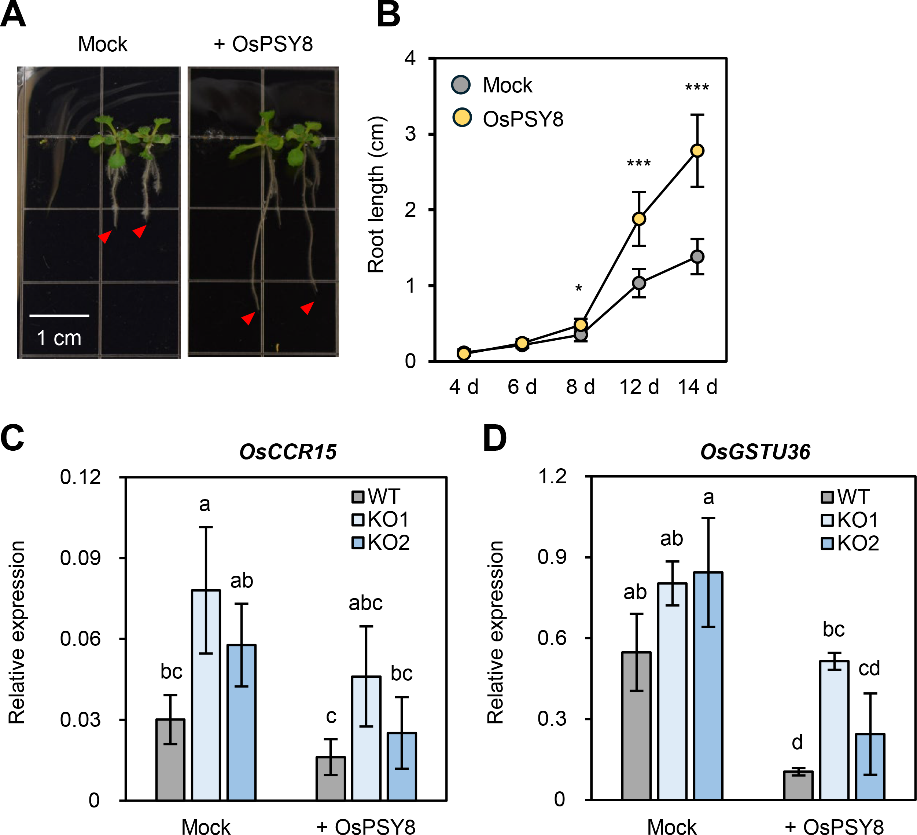

**Fig S5. Synthetic OsPSY8 peptide partially rescues elevated *OsCCR15* and *OsGSTU36* expression in KO roots**. (A) Representative images of 14-d-old *Arabidopsis tpst* seedlings treated with 500 nM synthetic sulfated OsPSY8 peptide. Mock, untreated control. Red arrowheads indicate root tips. (B) Root length of *tpst* seedlings treated with mock or OsPSY8 peptide was measured at 4, 6, 8, 12, and 14 days after germination. Data are mean ± SD (n = 6).  Asterisks indicate statistically significant differences from the mock control (**P* < 0.05, ****P* < 0.001; two-tailed Student's *t*-test). ns, not significant. (C,D) Expression level of *OsCCR15* (C) and *OsGSTU36* (D) in roots of WT, KO1, and KO2 plants under mock treatment and 2 h after treatment with 500 nM synthetic OsPSY8 peptide. Relative expression was quantified by RT-qPCR as the transcript level of each target gene normalized to rice *OsUBQ5* using the 2^−ΔΔCT^ method. Data are mean ± SD (n = 3 biological replicates), each replicate consisting of three plants. Statistical analyses were conducted using one-way ANOVA followed by Tukey's honestly significant difference (HSD) post hoc test for multiple comparisons, with different letters indicating significant differences among groups (*P* < 0.05).

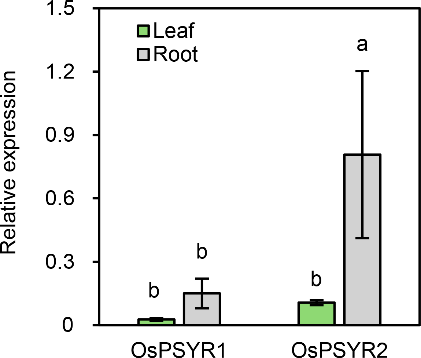

**Fig S6. Spatial expression of putative *OsPSYR* genes in leaves and roots of 9-d-old hydroponically grown rice seedlings.** Relative expression was quantified by RT-qPCR as the transcript level of each target gene, normalized to rice *UBIQUITIN5* (*OsUBQ5*) using the 2^−ΔΔCT^ method. Data represent mean ± SD (n = 3 biological replicates), each replicate consisting of three plants. Statistical analyses were conducted using one-way ANOVA followed by Tukey's honestly significant difference (HSD) post hoc test for multiple comparisons, with different letters indicating significant differences among groups (*P* < 0.05).

Tables

Table S1. Primers used in this study.

| **A. Primers for the generation and verification of overexpression plants** | | |
| --- | --- | --- |
| Primer name | Forward primer (5’→3’) | Reverse primer (5’→3’) |
| *OsPSY5*-OX | CACCATGGAGAGAGTTCCTGCAG | GCTGCTCAGTTTCTCCTCTCCTGC |
| *OsPSY8*-OX | CACCATGGCGCAGCAGCAGC | TCACGGATGCGGGTTGTGGC |
| transgene detection | GATGTAGGAGGGCGTGGATA | GATGTTGGCGACCTCGTATT |
| **B. Primers for the CRISPR/Cas9 vector construction and editing detection** | | |
| *ospsy8-1*_gRNA | TAGGTCTCCTCGATTGGAATGGTTTTAGAGCTAGAA | ATGGTCTCATCGAGATTCGCTTGCACCAGCCGGGAA |
| *ospsy8-2*_gRNA | TAGGTCTCCCATCAACGACTAGTTTTAGAGCTAGAA | ATGGTCTCAGATGGACTCCATTGCACCAGCCGGGAA |
| *ospsy5_gRNA* | TAGGTCTCCCACCTTGAGGGCGTTTTAGAGCTAGAA | TAGGTCTCCCACCTTGAGGGCGTTTTAGAGCTAGAA |
| Cas9_detection | CAGAAGAAGATACACCAGAC | GAACAGGCCATTCTTCTTCTC |
| editing_detection *_ospsy5* | ATGGAGAGAGTTCCTGCAGTT | GGTACAGGTACAAAGTCTG |
| editing*_*detection __*ospsy8* | CAGGTAGTACTCTGTTGCGC | GACGAGTGGTCAAACGTTAC |
| **C. Primers for the generation of GUS transgenic plants** | | |
| *pOsPSY5* | CTAGTTGGCTTCTTGAACTG | CTTGAAGTACTTTTCTAC |
| *pOsPSY8* | CACCTTGATGATGATCTCGTC | CACATTCTCCTCCTCTTCCTC |
| **D. Primers for RT-qPCR** | | |
| *OsPSY1* (Os05g0487100) | CGTGATCTCGTCGCCGTCTTG | TGGAGACGACAGGTGTTGCTG |
| *OsPSY2* (Os05g0487300) | TCTGTCCTGCCTCCTCCTC | CTGGAGAGAGGTTCAGAGACGG |
| *OsPSY3* (Os01g0276900) | TGCTCGGTGCCTTTCATCCTG | GACCGGCAGCTCTCCCTCTA |
| *OsPSY4* (Os01g0815400) | GCGTGTCCTTGAGGAACCAC | TTCGTCGAGGTCTCGCTCTTC |
| *OsPSY5* (Os11g0600600) | TGCAGTTCATCTGGCAGTGGT | TCCTCTGCTAGAGACTGGAGTTGC |
| *OsPSY6* (Os07g0631300) | GGGTGTTGGTGTTTCAGGTTCAGC | TCATGTGCCGGGTGCCTTG |
| *OsPSY7* (Os05g0542300) | ATGCCATCAGTTTCAGTTGCATCC | GGTATATGCTGTCTCAGCTTGCC |
| *OsPSY8* (Os01g0264400) | GACTCCGCTAGGACAGGAG | GGGTTGTGGCGGTTGTT |
| *OsUBQ5* (Os01g0328400) | ACCACTTCGACCGCCACTACT | ACGCCTAAGCCTGCTGGTT |
| *OsWRKY24* (Os01g0826400) | GACGACGAGATCAGAGTTGG | GTCGCTCATGGTTTGGACGAC |
| *OsGSTU36 (Os01g0949800)* | GGGCTCACTTCATCGAACAC | GAAGCGCCAGGTTCTCCTTC |
| *OsCCR15* (Os09g0491788) | CTTCTAGTGTACGACAAGGC | CGTAGTCATATCCACATCAACC |
| **E. Primers for dual-luciferase assay** | | |
| *pOsPSY8* _BamHI(F) and SalI(R) | GGATCCCACCTTGATGATGATCTCGTC | GTCGACCACATTCTCCTCCTCTTC |
| *OsWRKY24* _BamHI(F) and KpnI(R) | GGATCCATGACAACCTCGTCGTCCG | GGTACCGTAGAGCGAGTTCTGG |
| **F. Primers used for ChIP assays** | | |
| *pOsPSY8*_ChIP_a | GTGTGGTCCAAAACCGATACACG | CAACGTTGGGTACAAGACACG |
| *pOsPSY8*_ChIP_b | GTGTGTCTAGCTCCGTAAGAGCAAG | GCAACACGTGTTAAACTAGCTCTTGC |
| *pOsPSY8*_ChIP_c | CAAAAGTCCAACACGACACCACAC | GCCGTTTAATTTCACCCGTGTCC |
| *pOsPSY8*_ChIP_d | GAATAGAGTAGTGCGCAGGGGTG | GACAGTAATTCCCCTCTCGTG |
| *pOsPSY8*_ChIP_e | CAGGCAGGCAACGAGAGGCTTTTC | GGCTGCTTCTCCTTCCTTTCTGC |
| *OsUBQ5* | ACCACTTCGACCGCCACTACT | ACGCCTAAGCCTGCTGGTT |

Dataset S1. Differentially expressed genes between WT (4h) and WT (0h) (log₂FC = WT(4h)/WT(0h)). log₂FC values were generated by the edgeR model, which accounts for library size differences across sample groups. CPM (counts per million) was calculated with edgeR and uses only samples in comparison to estimate library size.

Dataset S2. Differentially expressed genes between KO (0h) and WT (0h) (log₂FC = KO(0h)/WT(0h)). log₂FC values were generated by the edgeR model, which accounts for library size differences across sample groups. CPM (counts per million) was calculated with edgeR and uses only samples in comparison to estimate library size.

Dataset S3. Differentially expressed genes between KO (4h) and WT (4h) (log₂FC = KO(4h)/WT(4h)). log₂FC values were generated by the edgeR model, which accounts for library size differences across sample groups. CPM (counts per million) was calculated with edgeR and uses only samples in comparison to estimate library size.

Dataset S4. Differentially expressed genes between OX (0h) and WT (0h) (log₂FC = OX(0h)/WT(0h)). log₂FC values were generated by the edgeR model, which accounts for library size differences across sample groups. CPM (counts per million) was calculated with edgeR and uses only samples in comparison to estimate library size.

Dataset S5. Differentially expressed genes between OX (4h) and WT (4h) (log₂FC = OX(4h)/WT(4h)). log₂FC values were generated by the edgeR model, which accounts for library size differences across sample groups. CPM (counts per million) was calculated with edgeR and uses only samples in comparison to estimate library size.
